# Human microglia extensively reconstitute in humanized BLT mice with human interleukin-34 transgene and support HIV-1 brain infection

**DOI:** 10.1101/2021.02.18.431677

**Authors:** Jianshui Zhang, Saroj Chandra Lohani, Yilun Cheng, Tao Wang, Lili Guo, Woong-Ki Kim, Santhi Gorantla, Qingsheng Li

## Abstract

Humanized bone marrow-liver-thymic (hu-BLT) mice develop a functional immune system in periphery but have a limited reconstitution of human myeloid cells, especially microglia, in CNS. Further, whether bone marrow derived hematopoietic stem and progenitor cells (HSPCs) can enter the brain and differentiate into microglia in adults remains controversial. To close these gaps, in this study we unambiguously demonstrated that human microglia in CNS were extensively reconstituted in adult NOG mice with human interleukin-34 transgene (hIL34 Tg) from circulating CD34+ HSPCs but no in hu-BLT NOG mice, providing strong evidence that human CD34+ HSPCs can enter adult brain and differentiate into microglia in CNS in the presence of hIL34. Further, the human microglia in the CNS of hu-BLT-hIL34 NOG mice robustly supported HIV-1 infection reenforcing the notion that microglia are the most important target cells of HIV-1 in CNS and demonstrating its great potential as an in vivo model for studying HIV-1 pathogenesis and evaluating curative therapeutics in both periphery and CNS compartments.

## Introduction

Microglia,the resident macrophages in the central nervous system (CNS), are the key resident immune cells to maintain neuronal homeostasis, defend against infections, and are associated with the pathogenesis of many neurodegenerative diseases (1-3). The ontology of adult brain microglia has been debated for a long time. The consensual view to date is that microglia in CNS is the seeding results of primitive hematomyeloid precursor cells from yolk sac and aorta-gonad-mesonephros region in early embryo life and proliferation in situ thereafter (4-8). However, multiple studies also showed that bone marrow derived cells can enter the brain and differentiate into microglia in adults (9-11).

Humanized mice (hu-mice) with a human immune system have been extensively used in investigating the ontology of immune cells, immunopathogenesis of human specific pathogens, and evaluating therapeutics as preclinical small animal models(12-15). Hu-mice generated by engrafting human CD34+ hematopoietic stem and progenitor cells (HSPCs) in neonatal life can reconstitute macrophages in the meninges and perivascular spaces, but rarely in the parenchyma of brain(16, 17). Similarly, humanized bone marrow-liver-thymic (hu-BLT) mice engrafted with human fetal liver and thymic tissues and HSPCs at adults develop a functional immune system in periphery but have a limited reconstitution of human myeloid cells, especially microglia, in CNS(18). We previously demonstrated that human interleukin-34 transgenic (hIL34-Tg) NOG mice engrafted intrahepatically with CD34+ HSPCs at birth significantly reconstituted microglial-like cells in the CNS (19). However, it remained unknown whether adult hIL34-Tg NOG mice could also reconstitute human microglia in CNS until this study. The hu-BLT mice are the best hu-mice in terms of human immune reconstitution, as they are engrafted with human fetal thymic tissues in addition to human liver tissues and liver derived CD34+ HSPCs where human T cells can receive differentiation and selection education in human thymic tissues(20, 21). This study has been poised to address two questions using the hu-BLT hIL34-Tg NOG (hu-BLT-hIL34) mouse model. First, we wanted to investigate whether human parenchymal microglia in the CNS could be reconstituted in adult mice from circulating myeloid precursor cells derived from CD34+ HSPCs in the hu-BLT mice. This is a fundamental question regarding the origin of human microglia in CNS at adults. The second question was to test the functionality of reconstituted human microglial cells in supporting HIV-1 replication in CNS and evaluate its utility as an in vivo model for investigating HIV-1 pathogenesis and evaluating curing therapeutics in both periphery and CNS compartments.

Using this unique system and by comparing two types of hu-BLT mice with and without hIL34 Tg received the same human donor tissues, we unambiguously demonstrated that human microglia in CNS can be extensively reconstituted in adult hIL34 Tg NOG mice but no in NOG mice, which provides strong evidence that human CD34+ HSPCs can enter adult brain and differentiate into microglia in the CNS in the presence of hIL34. Further, the human microglia in the CNS of hu-BLT-hIL34 mice robustly supported HIV-1 infection, which reenforced the notion that microglia are the most important target cells of HIV-1 in CNS and demonstrated its great potential as an in vivo model for studying HIV-1 pathogenesis and evaluating curative therapeutics in both periphery and CNS compartments.

## Method

### Ethics

All methods associated with animals described in this study were conducted in accordance with the Institutional Animal Care and Research Committee (IACUC) approved protocols at the University of Nebraska-Lincoln (UNL) and University of Nebraska Medical Center (UNMC).

### hIL34-Tg NOG and NOG mice

NOG (NOD.Cg-*Prkdc*^*scid*^ *Il2rg*^*tm1Sug*^/JicTac) mice of 6-8-week-old were purchased from Taconic Biosciences (Rensselaer, NY 12144, United States) and housed at the UNL Life Sciences Annex under specific-pathogen-free conditions. The hIL34-Tg NOG mice were bred at UNMC by pairing hIL34-Tg mouse with NOG mouse of opposite gender. The offspring were genotyped after three weeks of age by obtaining DNA from the tail snipping. Genotyping was performed for hIL34 as described previously using real-time polymerase chain reaction(19). Mice positive for hIL34 were transferred to UNL to generate hu-BLT mice.

### Generation of hu-BLT-hIL34 and hu-BLT Mice

To investigate human microglia reconstitution in the CNS of hu-BLT-hIL34 mice, both hu-BLT-hIL34 and hu-BLT mice were generated from adult hIL34-Tg NOG and NOG mice as we previously reported(22, 23) (Fig 1A). Briefly, 6-to-8-week-old adult hIL34-Tg NOG mice and NOG mice received sublethal irradiation at the dose of 12 cGy/gram of body weight with an RS2000 X-ray irradiator (Rad Source Technologies). Mice were surgically engrafted with a sandwich of two pieces of human fetal liver and one piece of thymic tissue fragments under the murine left renal capsules, of which human fetal livers and thymus tissues were procured from the Advanced Bioscience Resources. Within 6 hours after surgery, a total of 2.3 × 10^5^ fetal liver derived CD34^+^ HSPCs in 200 ul volume was injected through tail vein. At 16 weeks post engraftment, the immune reconstitution in the peripheral blood was assessed using flow cytometry as described below.

**Fig. 1.**
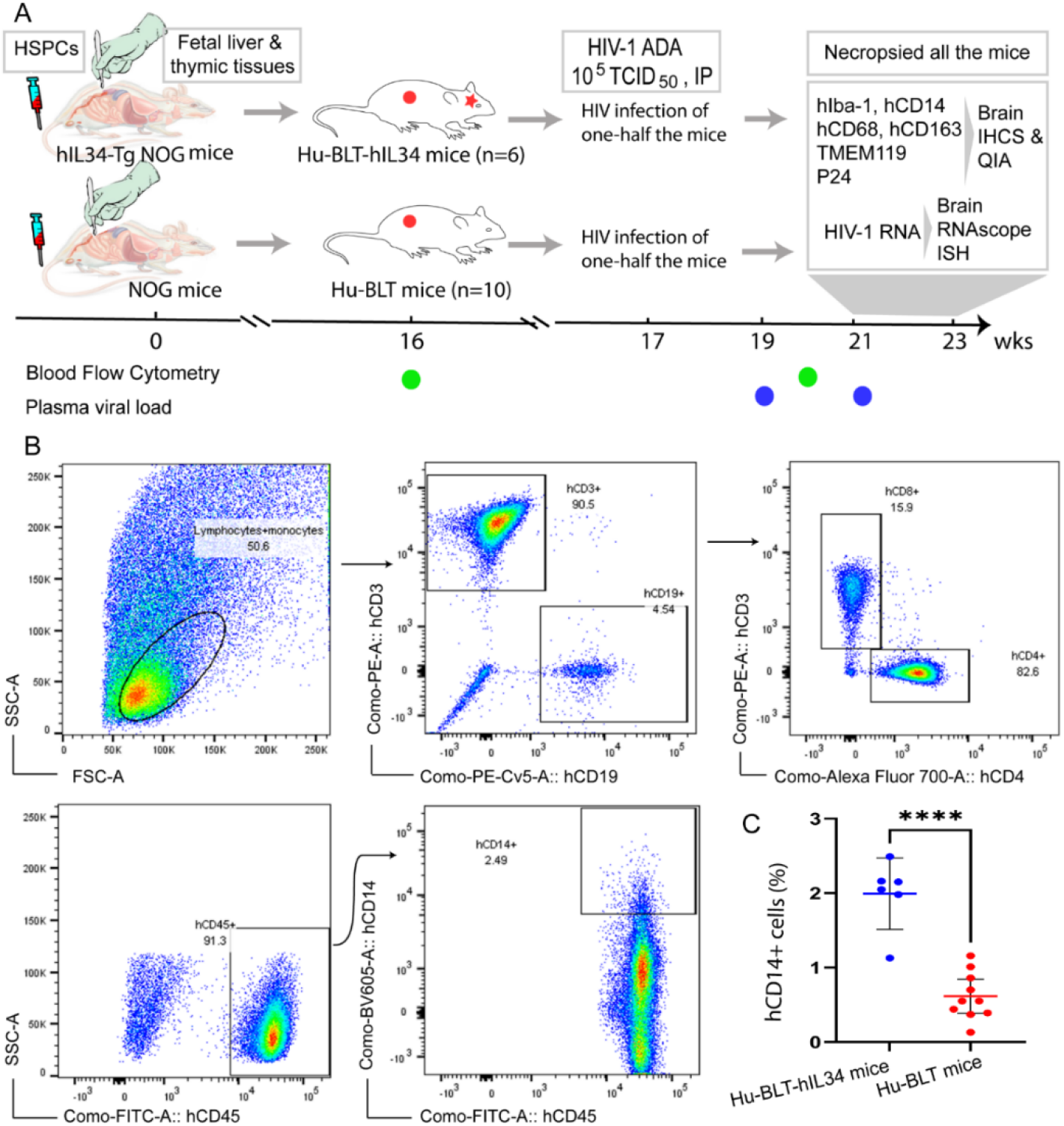
Experimental timeline and Flow Cytometric evaluation of human immune cells reconstitution in the peripheral blood of hu-BLT-hIL34 and hu-BLT mice. (**A**) Experimental design and timeline for the humanization of hIL34 transgenic mice and NOG mice by engrafting with human CD34+ hematopoietic stem and progenitor cells (HCSPs) and human fetal liver and thymic tissues, HIV-1 Ada infection of some mice in each group, and the evaluation of human macrophages and microglia reconstitution and HIV-1 infection in the CNS of euthanized mice using immunohistochemical staining (IHCS) and RNAscope in situ hybridization (ISH). (**B**) The representative flow cytometric dot plots of the peripheral blood mononuclear cells of a hu-BLT-hIL34 mouse (#1703) at the 4 months after transplantation, which were gated with hCD45+, hCD3+, hCD9+, and hCD14+ cells. (**C**) The percentage of hCD14+ myeloid cells in the peripheral blood of hu-BLT-hIL34 mice is significantly higher than that hu-BLT mice at the 4 months post transplantation. Each symbol represents an individual mouse, hu-BLT-hIL34 mice n=6, hu-BLT mice n=10; **** means p<0.0001.

### Human brain tissues

To compare the human microglia from hu-BLT-hIL34 mice with humans, ethically sourced human autopsy cerebral cortex tissues from a deidentified individual of HIV-1 negative with no registered medical complications were obtained from the NIH Neuro BioBank (https://neurobiobank.nih.gov/).

### Multicolor Flow Cytometry

Multicolor flow cytometry was conducted as previously reported (22, 23) for assessing the human immune reconstitution of peripheral blood mononuclear cells (PBMCs) at 16- and 20-weeks post transplantation. Briefly, peripheral blood was collected from a great saphenous vein into the ethylenediaminetetraacetic acid (EDTA)-containing vial (BD Microtainer, Franklin Lakes, NJ, USA). Red blood cells were lysed using fluorescence activated cell sorting (FACS) lysing solution (BD Biosciences, USA). The PBMCs were resuspended in a FACS buffer (2% FBS in phosphate-buffered saline) and incubated with a cocktail of following monoclonal antibodies against human immune cell markers at 4 °C for 30 min: anti-hCD45-fluorescein isothiocyanate (FITC), anti-hCD3-Phycoerythrin (PE), anti-hCD19-Phycoerythrin-Cyanin 5 (PE-Cy5), anti-hCD4-Alexa Fluor 700 (AF700), anti-hCD8-allophycocyanin-Cyanin 7(APC-Cy7) and anti-hCD14-Brilliant Violet 605 (BV605). All above antibodies and isotype control antibodies were obtained from BD Biosciences, USA. The stained cells were washed with FACS buffer and fixed with 4% paraformaldehyde. Raw data were acquired by FACSAria III (BD Biosciences) and analyzed with FlowJo version 7.6.4 (TreeStar). Statistical analysis was performed using GraphPad Prism 7 (GraphPad Software). Two groups of female hu-BLT-hIL34 (n=6) and hu-BLT mice (n=10) with good human immune cell reconstitutions in the peripheral blood (Table 1) were selected for this study.

### HIV-1 infection and measurement of HIV-1 plasma viral loads

To investigate the infectivity of HIV-1 in parenchymal human microglial cells in the CNS, 4 mice (# 1703, 1705, 1708 and 1709) of the hu-BLT-hIL34 mice group and 5 mice (# 1720, 1723, 1724, 1726, and 1728) in the hu-BLT mice group were randomly selected (Table 1) and intraperitoneally inoculated with the 10^5^ tissue culture infectious dose 50 (TCID50) of macrophage-tropic HIV-1 ADA in 200μl volume (obtained through the NIH AIDS Reagent Program, Division of AIDS, NIAID, NIH: HIV-1 ADA Virus from Dr. Howard Gendelman). At 2- and 4-weeks post HIV-1 inoculation, HIV-1 plasma viral load (pVL) in copies/ml was determined by real-time RT-PCR using our previously published protocol(24). Briefly, viral RNA was extracted from the plasma using QIAamp ViralRNA minikit (Qiagen) as recommended by the manufacturer and quantified using C1000 ThermalCycler and the CFX96 Real-Time system (Bio-Rad). For 20 µl qRT-PCR, 5 µl of extracted viral RNA, TaqMan Fast Virus 1-Step master mix (Life Technologies) and following primers and probe combination (IDT, USA) were used: Forward Primer, GCCTCAATAAAGCTTGCCTTGA; Reverse Primer, GGGCGCCACTGCTAGAGA; Probe, /56-FAM/CCAGAGTCA/ZEN/CACAACAGACGGGCACA/3IABkFQ/.

### Euthanasia of all the mice for evaluating human myeloid cell reconstitution and HIV-1 infection in the CNS

After two consecutive positive results of HIV-1 pVL at 2- and 4-weeks post HIV-1 inoculation, which is equivalent to the 21- to 23-weeks post transplantation, all the mice, including HIV-1 infected and non-inoculated subgroups from the hu-BLT-hIL34 and hu-BLT mice groups, were euthanized for analyzing human myeloid cell reconstitutions and HIV-1 infections in the CNS (Fig. 1A). Whole brain was dissected out during necropsy and sliced coronally into 5 parts at 4 mm interval using a young mouse brain slicer (Cat# BSMYS001-1, Zivic Instruments, Pittsburgh, PA, USA). The brain tissues and other tissues including spinal cord, spleen, lymph node, jejunum and ileum were collected and fixed in SafeFix™ II (Cat# 042600, Fisher Scientific, USA) at room temperature for 6 hours and embedded in paraffin.

### Immunohistochemical staining (IHCS)

To evaluate the phenotype, morphology and distribution of the reconstituted human myeloid cells in the CNS, IHCS was conducted by following our previously published protocol with slight modifications (25). Briefly, antigen retrieval of 6-μm thick tissue sections were performed in 0.1mM citrate buffer (PH 6.0) by heating at 98°C for 15 mins. The following primary antibodies were used for detecting macrophage/microglial cells: rabbit monoclonal-antibody (mAb) to human ionized calcium-binding adaptor molecule 1 (hIba-1, EPR6136-2 clone, Cat# ab221933, 1:500; Abcam, USA), rabbit polyclonal antibody to human TMEM119-C-terminal (hTMEM119, Cat# ab185333, 1:500; Abcam, USA), rabbit mAb to human CD14 (hCD14, EPR3653 clone, Cat. # 133335, 1:1000; Abcam, USA), mouse mAb to human CD68 (hCD68, PG-M1 clone, Cat# MS-1808-S1, 1:100; Thermoscientific, USA), mouse mAb to human CD163 (hCD163, 10D6 clone, Cat# NCL-L-CD163, 1:100; Leica Biosystems, USA). For HIV-1 detection, mouse mAb to HIV-1 gag p24 (Kal-1 clone, Cat# M0857, 1:10, Dako, USA) was used. Mouse or rabbit IgG isotype control antibodies were used as negative control. Dako EnVision+ system-HRP labelled polymer anti-rabbit kit (Code K4002, Dako, USA) or anti-mouse kit (Code K4000, Dako, USA), and the Betazoid DAB Chromogen Kit (Cat# BDB2004, BioCare Medical, USA) were used for signal detection and visualization. The cell nuclei were counterstained with Mayer’s hematoxylin. The stained tissue sections after completion of IHCS were digitized with Aperio CS2 Scanscope and the quantitative image analysis of positive cells was conducted using a positive pixel count algorithm in Aperio’s Spectrum Plus analysis program (version 9.1; Aperio ePathology Solutions) as previously reported(26).

### HIV-1 viral RNA detection using RNAscope in situ hybridization (ISH)

HIV-1 viral RNA (vRNA) in the brain tissues were detected using RNAscope ISH according to our previously published protocol (26). Briefly, HIV-1 antisense probes of RNAscope® ISH probe-V-HIV1-clade B (Cat#416111) and RNAscope® 2.5 HD assay-Red kit were used. The RNAscope® probe-DapB (Cat# 310043) was used as a negative control. All the reagents above were purchased from the Advanced Cell Diagnostics, Inc.

### Combined RNAscope ISH with IHCS

To determine the cell types of vRNA+ cells in the CNS, a combined RNAscope ISH and IHCS method was used as reported (26). Briefly, after the completion of RNAscope ISH for HIV-1 vRNA and digitization of the whole tissue section, the slides were soaked in xylene overnight to remove the coverslips and the tissue sections were rehydrated and subjected to IHCS using the rabbit mAb to hIba-1 ((EPR6136-2 clone, Cat# ab221933, 1:500; Abcam, USA)) as the primary antibody as described in the IHCS section above. Rabbit IgG isotype control antibody was used as negative control.

## Results

### Human myeloid cell reconstitution in the CNS

CNS contains four types of macrophages: parenchymal microglia and nonparenchymal perivascular macrophages, meningeal macrophages as well as choroid plexus macrophages (2, 8). First, we used antibodies to hIba-1 and hCD14, sensitive markers of macrophages and microglial cells (27-30), to evaluate human myeloid cells reconstitution in the CNS of hu-BLT-hIL34 and hu-BLT mice. There were extensively reconstitutions of human myeloid cells in the CNS parenchyma of hu-BLT-hIL34 mice as indicated by hIba-1+ or hCD14+ positive cells using IHCS in the brain and spinal cord (Fig. 2 & 3). As shown in a representative whole brain tissue section from the third coronary slice (Fig. 2C), hIba-1+ cells in the brain parenchyma were extensively reconstituted across multiple regions of the brain. Fig. 2A and 2D respectively highlighted the cerebral cortex and hippocampus boxed regions from the Fig. 2C; in turn, Fig. 2B and 2E respectively further highlighted the boxed regions from the Fig. 2A and 2D at a higher magnification. The hIba-1+ cells are numerous, morphologically ramified, distributed in brain parenchyma. Consistent with the extensive reconstitution of parenchymal human myeloid cells in the CNS revealed by hIba-1+ cells, there were also abundant hCD14+ myeloid cells as shown in a representative whole brain tissue section (2G) of a hu-BLT-hIL34 mouse (#1708). The blue and red boxed regions of cerebral cortex and hippocampus in the Fig. 2G was respectively highlighted at a higher magnification in the Fig. 2F and 2H. The hCD14+ cells are again morphologically ramified and mainly distributed in brain parenchyma. Consistent with the abundant reconstitutions of human myeloid cells in the brain, we also observed the similar reconstitutions in the spinal cord **(Fig. 3**). The upper panel of Fig. 3 shows representative whole spinal cord tissue sections (**A, C**) and the highlighted images (**B, D**) form the corresponding boxed regions from hu-BLT-hIl34 mouse #1708 that were stained immunohistochemically for hIba-1+ or hCD14+ cells (brown) and counterstained with hematoxylin for cell nuclei. There were abundant hIba-1+ cells (**A** & **B**) and hCD4+ cells (**C** & **D**) in the spinal cord of hu-BLT-hIl34 mouse. We thus concluded that hu-BLT-hIL34 mice extensively reconstituted parenchymal human myeloid cells in the CNS, including cerebral cortex, hippocampus, thalamus hypothalamus, striatum, amygdala, spinal cord, and cerebellum (Supplemental Fig. 1). In contrast and as expected, there were limited reconstituted human myeloid cells in the parenchyma of the CNS of the hu-BLT mice based on hIba-1 (Fig. 3G-K). As shown in a representative whole brain tissue section from the third coronal slice (Fig. 3I, left upper), Iba-1+ cells were rare in all the parenchyma of cerebral brain (Fig 3I). Similarly, hCD14+ cells were also rare in the parenchyma of hu-BLT mice (Fig. 3 E-F). We next compare the reconstitution of other three types of nonparenchymal perivascular macrophages, meningeal macrophages, and choroid plexus macrophages in the hu-BLT-hIL34 and hu-BLT mice. The subset of hIba-1+ or hCD14+ cells from hu-BLT-hIL34 mice also expressed hCD68 and hCD163. As indicated by hIba-1, hCD14, hCD163 expression and anatomic distribution, hu-BLT-hIL34 mice also have a better reconstitution of meningeal macrophages (MM) and perivascular macrophages (PVM) (Supplemental Fig. 2). than hu-BLT mice (Fig 3 G-K).

**Fig. 2.**
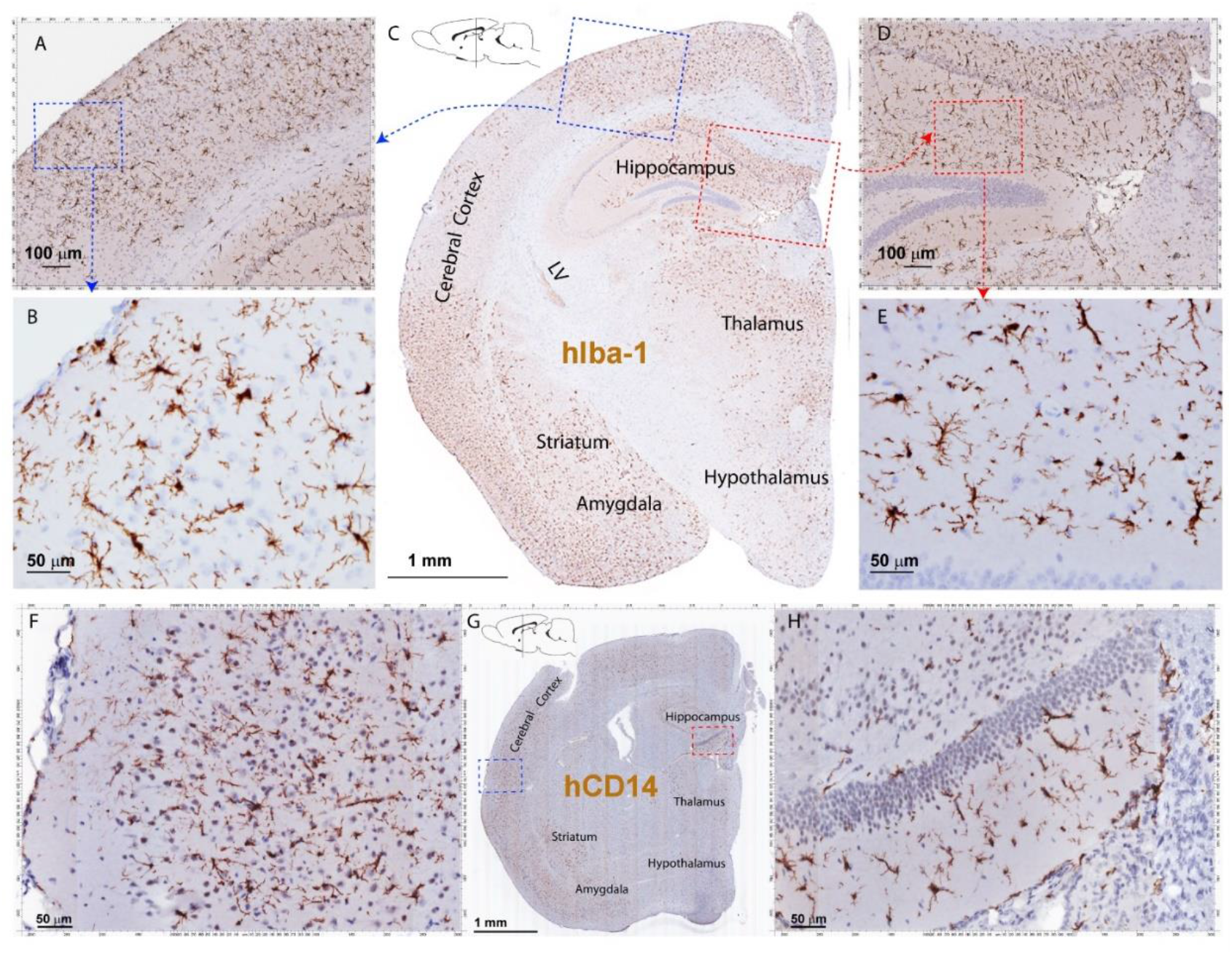
The hIba-1+ and hCD14+ myeloid cell reconstitution in the brain of hu-BLT-hIL-34 BLT mice. The upper panel (**A-E**) shows a representative whole brain tissue section from the third coronal brain slice of a hu-BLT-hIL34 mouse (#1703) that was stained immunohistochemically for hIba-1+ (brown) and counterstained with hematoxylin. LV stands for lateral ventricle. The blue and red boxed regions in the Fig. **C** were highlighted at a higher magnification (**A, D)**. In turn, the blue and red boxed regions of the Fig A and B were further highlighted (**B, E)**. The lower panel (**F-H**) shows a representative whole brain tissue section (**G)** of a hu-BLT-hIL34 mouse (#1708) that was stained immunohistochemically for hCD14+ cells (brown). The blue and red boxed regions was highlighted at a higher magnification (**F, H**). The hIba-1+ or hCD14+ cells are morphologically ramified and mainly distributed in brain parenchyma.

**Fig. 3.**
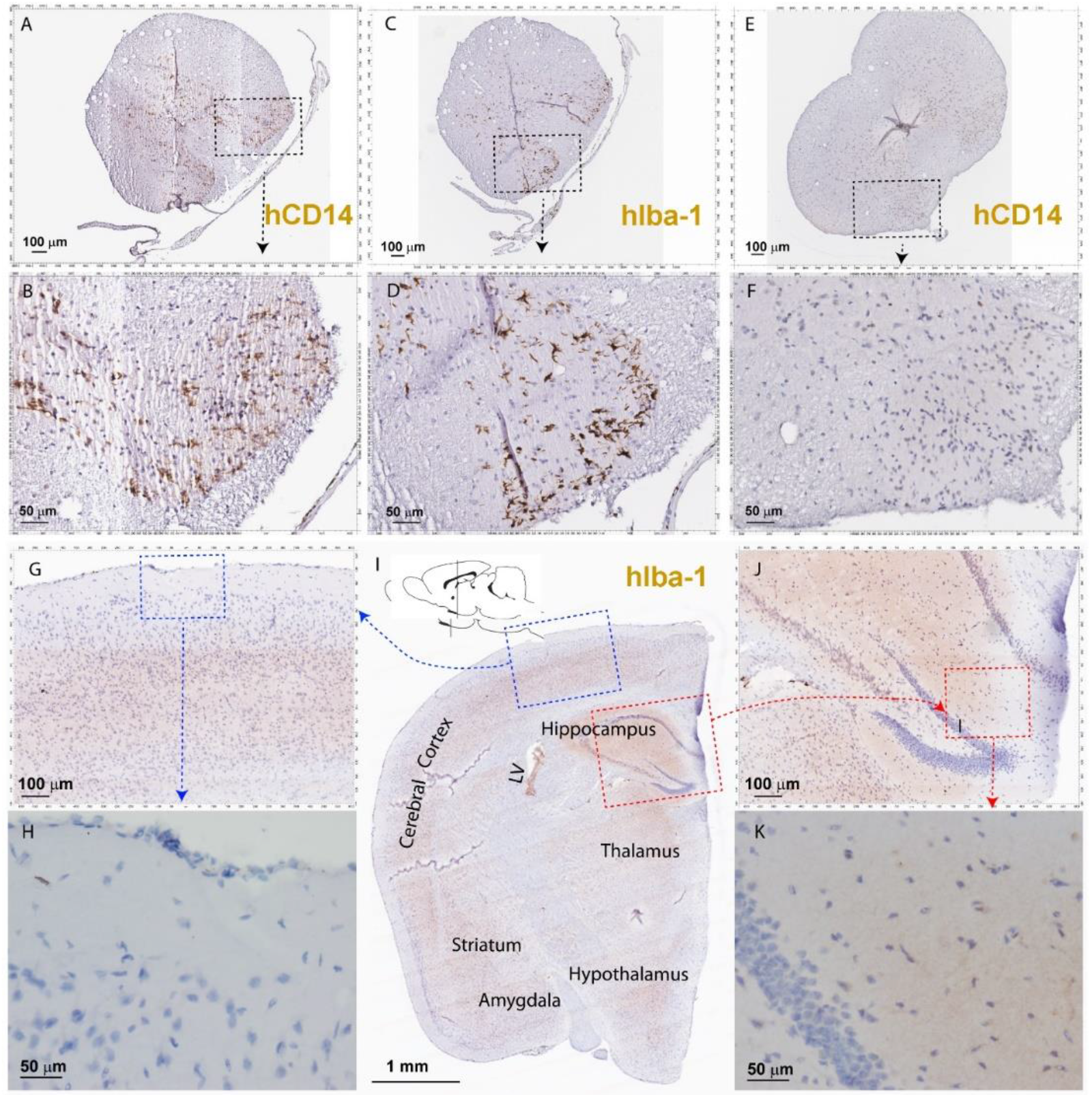
The hIba-1+ and hCD14+ myeloid cell reconstitutions in the spinal cord tissues of hu-BLT-hIL-34 BLT mice and rare myeloid cells in the brain and spinal cord parenchyma of hu-BLT mice. The upper panel (**A-F**) shows representative whole spinal cord tissue sections, and the highlighted images form the corresponding boxed regions from hu-BLT-hIl34 mouse #1708 (**A-D**) and hu-BLT mouse #1717 (**E-F**) that were stained immunohistochemically for hIba-1+ or hCD14+ cells (brown) and counterstained with hematoxylin. There were abundant hIba-1+ cells (**A** & **B**) and hCD4+ cells (**C** & **D**) in the spinal cord of hu-BLT-hIl34 mouse, where rare myeloid cells of hCD14+ cells (**E** & **F**) from the spinal cord tissues of the hu-BLT mouse. The lower panel (**G-K**) shows there were rare hIba-1+ myeloid cells in a representative whole brain tissue section from the third coronal slice of a hu-BLT mouse (#1717). LV stands for lateral ventricle. The blue and red boxed regions in the Fig. I were highlighted at a higher magnification (**G, J)**. In turn, the blue and red boxed regions of the Fig G and J was highlighted (**H & K)**.

### The myeloid cells in brain parenchyma expressed microglia-specific marker TMEM119

To distinguish parenchymal microglia from macrophages, we conducted IHCS using microglial-specific marker hTMEM119 (31, 32). There were extensively reconstitutions of hTMEM119+ cells in the CNS parenchyma of hu-BLT-hIL34 mice (Fig. 4). As shown in a representative whole brain tissue section from the third coronal slice (Fig. 4C), hTMEM1191+ cells in the brain parenchyma were extensively reconstituted across multiple regions of the brain. The Fig. 4A and 4D respectively highlighted the boxed regions of the cerebral cortex and hippocampus from the Fig. 4C; in turn Fig. 4B and 4E respectively further highlighted the boxed regions from the Fig. 4A and 4D at a higher magnification. The hTMEM119+ cells are morphologically ramified and distributed in parenchyma. We quantified TMEM119 + cells in the cerebral cortex of the hu-BTL-hIL34 mice (n=6) and found there were 304.08±131.93 (mean ± SD) hTEME119+ microglial cells/mm2, whereas there were absent of these cells in the hu-BLT mice. We further compared hTMEM1109 + cells in the cerebral cortex of an HIV-1 non-infected individual, who had no registered medical complications at NIH Neuro Biobank (Fig4. G-I). We found that the frequency, morphology, and distribution of hTMEM119+ cells in hu-BLT-hIL34 mice are similar to that person (Fig 4. G-I). We thus concluded that hu-BLT-hIL34 mice extensively reconstituted human microglia in the CNS.

**Fig. 4.**
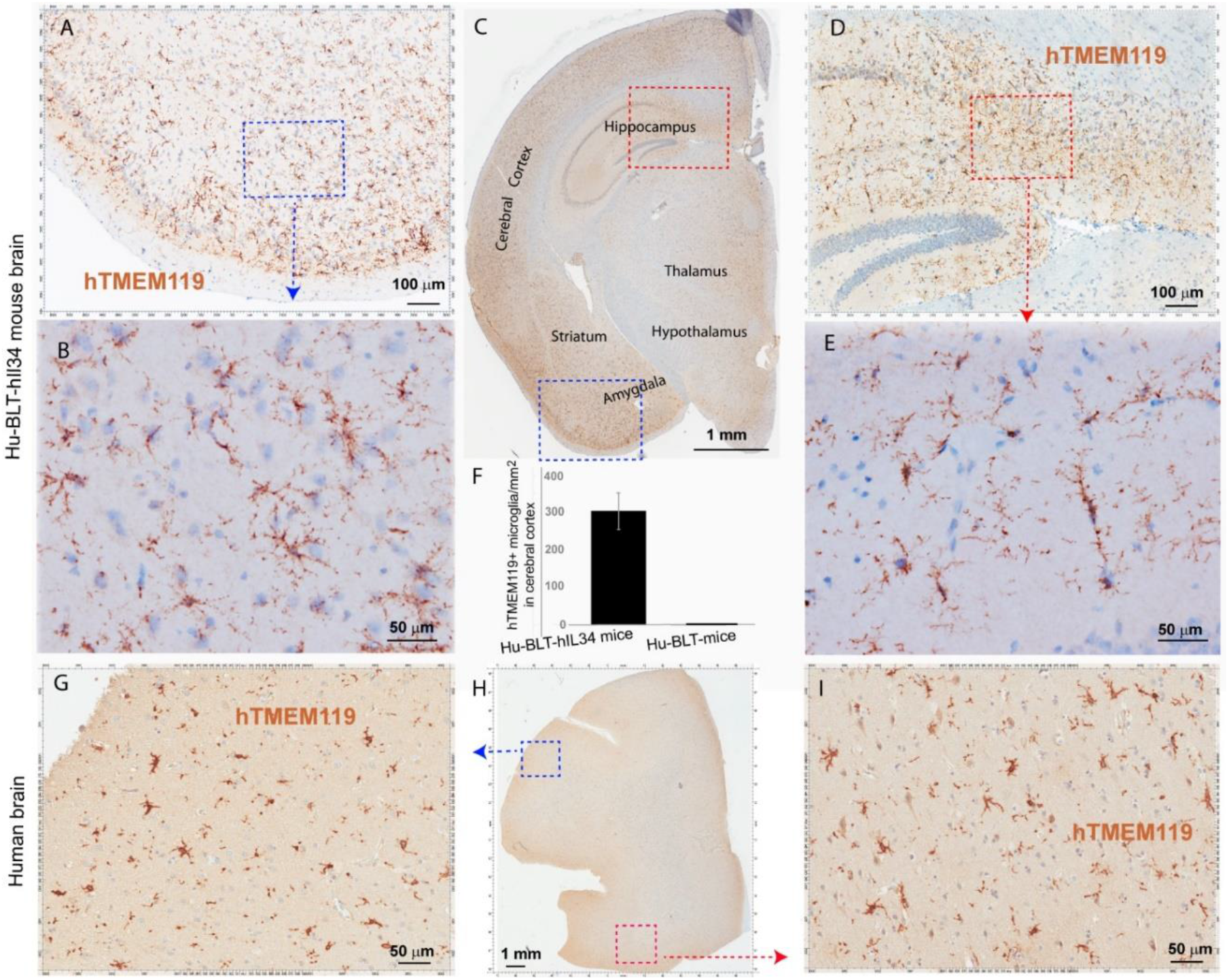
Microglia-specific marker TMEM119+ microglia in the brains of hu-BLT-hIL-34 mice and humans. A representative whole brain tissue section (**C**) from the third coronal brain slice of a hu-BLT-hIL34 mouse (#1703) that was stained immunohistochemically for hTMEM119+ cells (brown) and counterstained with hematoxylin. LV stands for lateral ventricle. The blue and red boxed regions in the Fig. C were highlighted at a higher magnification (**A** & **D)**. In turn, the blue and red boxed regions of the Fig A & D were highlighted (**B** & **E)**. The hTMEME119+ cells (brown) are morphologically ramified and distributed in brain parenchyma. The histogram of quantitative images analysis of TMEM119 + cells in the cerebral cortex of the hu-BTL-hIL34 mice (**F**) (304.08±131.93 hTEME119+ cells/mm2, n=6), where there were absent of these cells in the hu-BLT mice. The lower panel (G-I) shows hTMEM119 + microglia in the cerebral cortex of a HIV-1 non-infected individual with no registered medical complications (Fig 4. G-I). The frequency, distribution, and morphology of TMEM119+ human microglia in hu-BLT-hIL34 mice are similar to this human individual.

### HIV-1 infection in the CNS of hu-BLT-hIL34 mice

To test the functionality of reconstituted human myeloid cells in the CNS of hu-BLT-hIL34 mice, we infected 4 animals from hu-BLT-hIL34 group and 5 animals from hu-BLT mice groups. At 4-6 weeks post HIV-1 infection, we euthanized all animals and detected abundant HIV-1 vRNA+ and p24+ cells in the brain of hu-BLT-hIL34 mice (Fig. 5) but rarely in hu-BLT mice (data not shown). As shown in a representative image in the Fig. 5, there are abundant HIV-1 RNA+ cells (red) in the brain tissues of hu-BLT-hIL34 mouse (#1708) detected using RNAscope ISH with HIV-1 clade B probe. Consistent with the results of HIV-1 VRNA+ cells, there were abundant HIV-1 p24+ cells detected using IHCS. We further defined the HIV-1 vRNA+ cells expressed as human myeloid cell marker hIba-1 (arrows) indicating the reconstituted human myeloid cells can support HIV-1 infection.

**Fig. 5.**
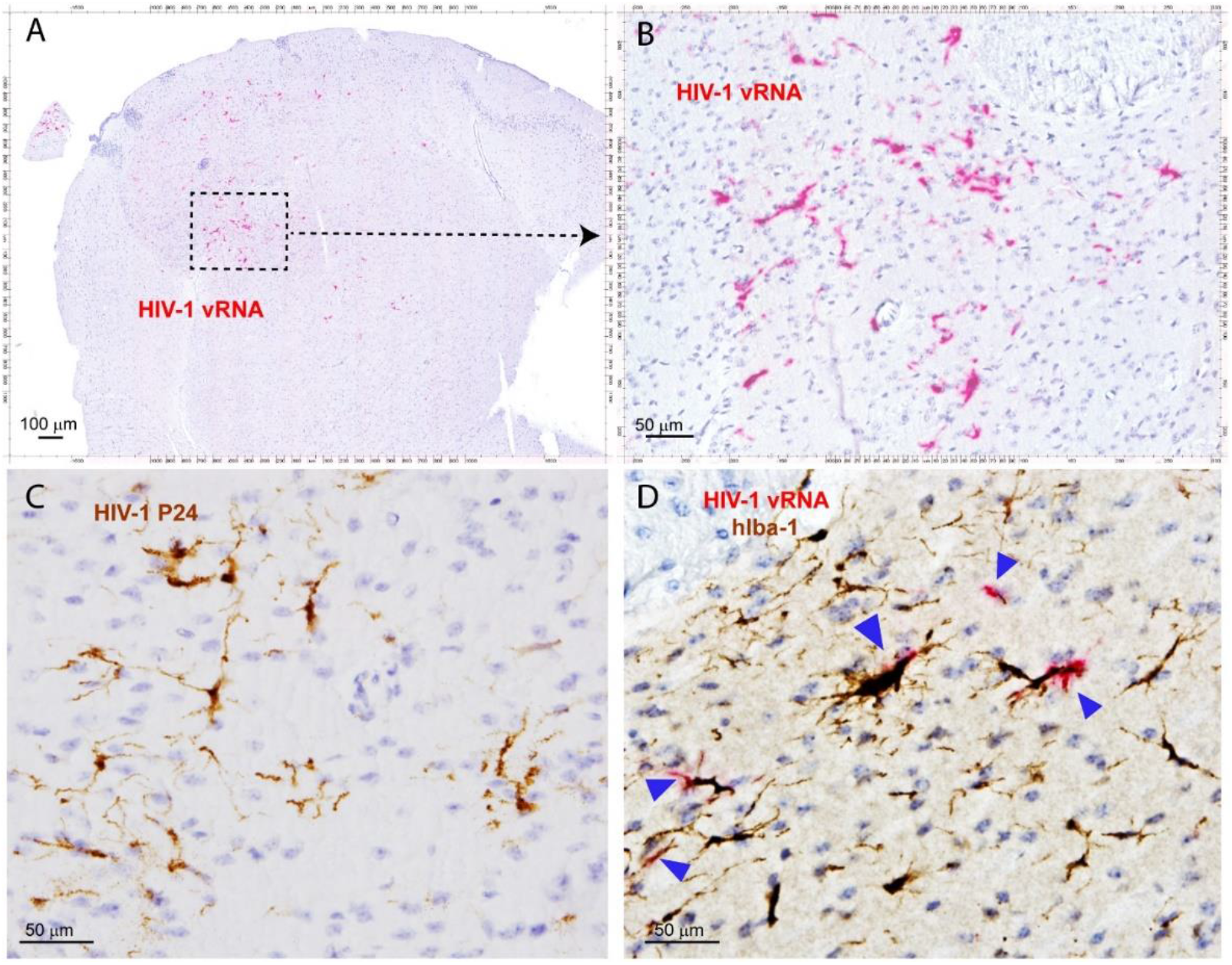
HIV-1 infection of human microglia of hu-BLT-hIL34 mice. **(A)** HIV-1 RNA+ cells (red) in a representative whole brain tissue section from a hu-BLT-hIL34 mouse (#1708) that was detected using RNAscope in situ hybridization with HIV-1 clade B probe and counterstained with hematoxylin. (**B**) Highlighted boxed region at a higher magnification from the Fig A showing HIV-1 RNA+ cells (red). (**C**) HIV-1 p24+ cells (brown) in the cerebral cortex of hu-BLT-hIL34 mouse (#1708) detected using IHCS detected. (**D**) Colocalization of HIV-1 vRNA and human myeloid cell marker hIba-1 (arrows) indicating the reconstituted human myeloid cells could support HIV-1 replication.

## Discussion

In this study, we demonstrated that human microglia can be extensively reconstituted in CNS from circulating human HSPCs in hu-BLT-hIL34 mice. We first used a battery of human myeloid cell markers, including hIBa-1, hCD14, hCD68 and hCD163, to evaluate human myeloid cell reconstitution in the CNS and found that human myeloid cells were extensively reconstituted and primarily localized in the brain parenchyma (Fig 2 & 3). We then used a human microglial specific marker, hTMEM119, to validate these reconstituted human myeloid cells in the brain parenchyma are mainly human microglia (Fig 4. A-F). Further, in comparison with hTMEM119 + microglia in the cerebral cortex of a HIV-1 non-infected individual with no registered medical complications (Fig 4. G-I), we found the frequency, distribution, and morphology of TMEM119+ human microglia in hu-BLT-hIL34 mice are similar to this person. Our data thus support the notion that human microglia at adults can be generated through human hematopoietic stem and progenitor cells (HSPC), which is consistent with the previous reports that bone marrow derived cells can enter brain to different into microglia at adults (9-11). In contrast, we did not observe human microglia reconstitution in the brain parenchymal of hu-BLT mice. The hu-BLT-hIL34 and hu-BLT mice are genetically identical and also received the same human donor transplant except the former had hIL34 knock-in, indicating that hIL34, a ligand of the colony stimulating factor-1 receptor, play an important role in myeloid and microglial cells development in CNS (19, 33). This study is unique in the following aspects. First human myeloid and microglia cells reconstitution in the CNS is a clear-cut result in this chimeric mouse and human model. Second, the adult mice engrafted with HSPCs extensively reconstituted human microglia in the CNS, to our knowledge this is the first report in this regard. In addition to comparing the parenchymal human microglia between hu-BLT-hIL34 and hu-BLT mice, we also observed that hu-BLT-hIL34 mice had a much better reconstitution of meningeal, perivascular, and choroid plexus macrophages than hu-BLT mice. We also would like to point that both hu-BLT-hIL34 and hu-BLT mice received sublethal irradiation, whether this irradiation facilitated HSPCs to gain an entry into the brain and whether without irradiation can also reconstitute the brain microglia in adult hIl34-NOG mice remains to be investigated.

Despite the importance of microglial cells, as a resident macrophage, in host immune response to brain infections and in the pathogenesis of neurodegenerative diseases, very limited small animal models are available to recapitulate diseases pathogenesis associated with human microglia. To that end, we infected hu-BLT-hIL34 mice and found that reconstituted human microglia can support HIV-1 infection in the CNS. Moreover, HIV-1 vRNA were localized in human myeloid cells in the brain, reenforcing that microglia cells are the most important subtract of HIV-1 infection in the CNS(26, 34). Using hu-BLT-hIL34 mice, it is now feasible to investigate interplay between human pathogens, such as HIV-1, with human immune system of periphery and CNS compartments. The hu-BLT-hIl34 mouse model reported here open a new avenue for investigating the pathogenesis of HIV-1 infection and purging HIV-1 latent reservoir in the CNS in additional to peripheral tissues.

## Acknowledgments

We would like to thank University of Nebraska—Lincoln Life Sciences Annex and their staff for their assistance.

