## Supplemental Table 1, Fig. 1 & 2 for "Human microglia extensively reconstitute in humanized BLT mice with human interleukin-34 transgene and support HIV-1 brain infection"

**Supplemental Table 1** HIV-1 infection of hu-BLT-hIL34 and hu-BLT mice

| Group | Sub-group | Mouse ID | PVL, 4 wks PI |
|---|---|---|---|
| Hu-BLT-hIL34 Mice | HIV-1 infection | 1703 | 8.45E+05 |
| | | 1705 | 1.09E+06 |
| | | 1708 | 4.56E+06 |
| | | 1709 | 1.36E +05 |
| | Un-infection | 1699 | n/a |
| | | 1707 | n/a |
| Hu-BLT Mice | HIV-1 infection | 1720 | 8.28E+05 |
| | | 1723 | 5.78E+05 |
| | | 1724 | 1.09E+06 |
| | | 1726 | 1.36E+06 |
| | | 1728 | 5.46E+06 |
| | Un-infection | 1717 | n/a |
| | | 1718 | n/a |
| | | 1721 | n/a |
| | | 1722 | n/a |
| | | 1729 | n/a |

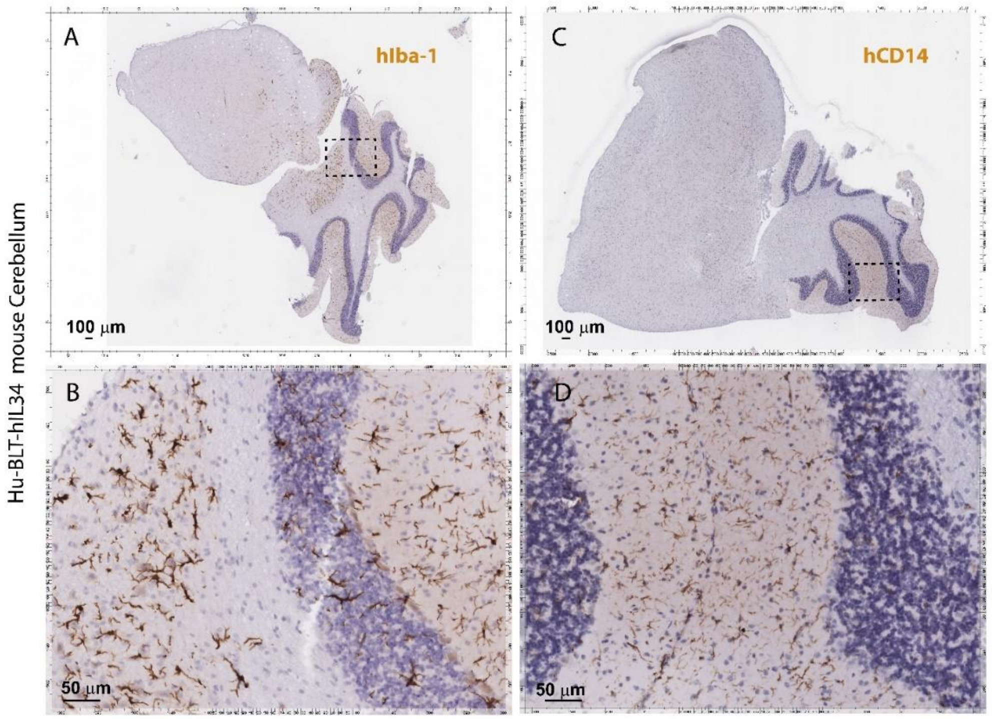


**Supplemental Fig 1. The hIba-1+ and hCD14+ myeloid cell reconstitutions in the cerebellum of hu-BLT-hIL-34**

**BLT mice.** Representative cerebellum tissue sections of hIba-1+ or hCD14+ cells from hu-BLT-hIl34 mice of #1703 respectively. There were abundant hIba-1+ cells (**A** & **B**) and hCD4+ cells (**C** & **D**)

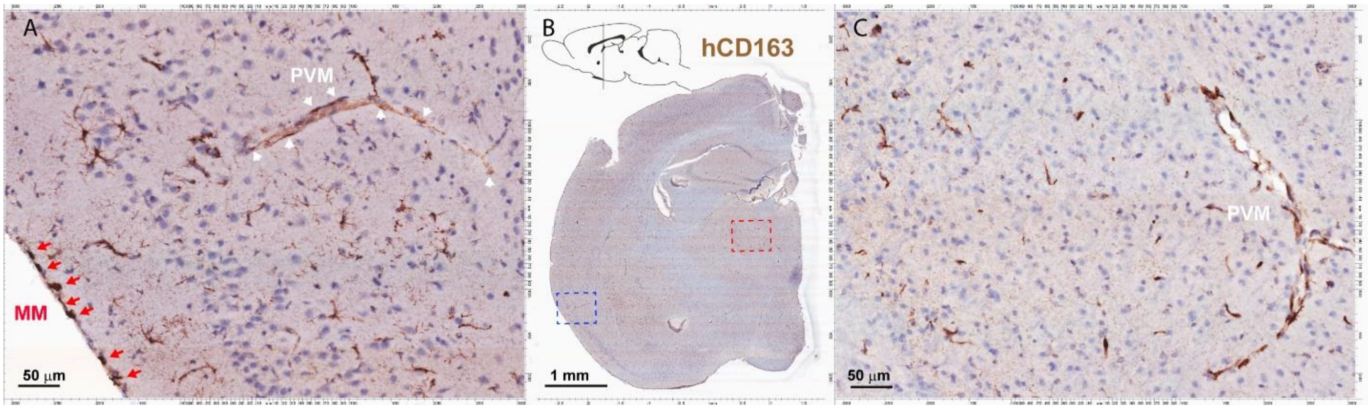


**Supplemental Fig 2. The hCD163+ myeloid cell reconstitutions in the brain of hu-BLT-hIL-34 BLT mice.** Representative brain tissue sections of hCD163+ cells from hu-BLT-hIl34 mouse (#1708). There were abundant hCD163+ meningeal macrophages (MM) and perivascular macrophages (PVM)
